# Typical hippocampal transcriptional response across estrous is dysregulated by *Cnih3* gene deletion

**DOI:** 10.1101/2022.04.11.487915

**Authors:** Bernard Mulvey, Hannah E. Frye, Tania Lintz, Stuart Fass, Eric Tycksen, Elliot C. Nelson, Jose A. Morón, Joseph D. Dougherty

**Affiliations:** Department of Genetics, Washington University School of Medicine, St. Louis, MO 63110, USA; Department of Psychiatry, Washington University School of Medicine, St. Louis, MO 63110, USA; Department of Anesthesiology, Washington University School of Medicine, St. Louis, MO 63110, USA; McDonnell Genome Institute, Washington University School of Medicine, St. Louis, MO 63110, USA

## Abstract

The hippocampus is a critical brain region for coordinating learning, memory, and behavior. In females, the estrous cycle alters these functions through steroid hormone activity, with well-characterized effects on cellular physiology and behavior. However, the molecular basis of these outcomes has not been systematically explored. Therefore, we profiled the transcriptome of dorsal hippocampi from female mice in each estrous cycle stage, and contrasted it with that of males. We identify only subtle sex differences in gene expression between the sexes on average, yet comparing males to individual estrous stages reveals up to thousands of genes deviating from male expression patterns at specific estrous stages. These estrous-responsive genes are especially enriched in gene markers of oligodendrocytes and the dentate gyrus, and in functional gene sets relating to estrogen response, potassium channels, and synaptic gene splicing. Next we profiled *Cnih3* knockouts across estrous to provide insight into their previously reported estrous-dependent phenotypes in hippocampal synaptic plasticity, composition, and learning and memory behaviors. Surprisingly, *Cnih3* knockouts showed far broader transcriptomic differences between estrous cycle stages and males. Moreover, *Cnih3* knockout drove subtle but extensive expression changes accenting sex differential expresssion at diestrus and estrus. Altogether, our profiling constitutes both a resource characterizing estrous-specific gene expression patterns in the adult hippocampus, which can provide insights into mechanisms of sex differential neuropsychiatric functions and dysfunctions, while also highlighting roles of *Cnih3* as a buffer against transcriptional effects of estrous and providing insights into the molecular mechanisms that may underlie estrous-dependent phenotypes with its loss.

## Introduction

Sex differences in brain and behavior are well-established factors in physiologic and pathologic processes ranging from reproduction and parenting to depression and addiction. Circulating sex hormones exert acute “activational” effects (Arnold, 2009) on cellular and behavioral phenotypes. Regular fluctuations in females across the human (menstrual) or rodent (estrous) cycle are known to continually alter the microstructure of the brain; for example, hippocampal synaptic density peaks when estradiol and progesterone are highest during proestrus (Woolley and McEwen, 1992), which has been shown to affect learning and memory (Frick et al., 2015a; Frick and Kim, 2018). Likewise, hormones influence the function of signaling within the brain and between tissues, as estrogens and progestins alter dopamine turnover, suppress GABA signaling (Del Río et al., 2018), and stimulate hypothalamic-pituitary-adrenal (HPA) axis-driven corticosteroid release by upregulating *CRH* (in contrast to its repression by androgens) (Bao et al., 2006; Lund et al., 2004).

Outwardly, the impact of sex has been observed in neuropsychiatric diseases and rodent models thereof, including addiction (Kuhn, 2015), potentially via the hippocampus (Kohtz and Aston-Jones, 2017). Indeed, it was recently observed that knockout (KO) of the opioid dependence-associated gene *Cnih3* (Nelson et al., 2016) has not only sex-, but estrous stage-specific effects (Frye et al., 2021) in learning and memory, supporting a role for acute sex hormonal effects on addiction-relevant brain circuitry related to learning and memory. Consistent with *Cnih3* modulating learning, KO showed severe memory deficits and corresponding synaptic changes, *but* only in female mice, and limited to particular phases of the estrous cycle (Frye et al., 2021). This suggested the surprising hypothesis that *Cnih3* is involved in buffering the female brain against effects of hormones on hippocampal learning, and thus loss of *Cnih3* results in these abnormal outcomes in a cycle-dependent manner.

While sex hormone receptors are thought to act primarily as transcriptional activators, the manner in which the estrous cycle alters brain gene expression, even in wildtype (WT) rodents, has only been characterized in a limited phases (Iqbal et al., 2020) or not yet directly contrasted to males (DiCarlo et al., 2017). Moreover, the brain functions of *Cnih3* are poorly understood, and are likely multifaceted. Thus, we sought to transcriptionally characterize the mouse hippocampus at each stage of the estrous cycle in both WT and *Cnih3* KO mice to clarify the molecular effects of KO and the transcriptional underpinnings of sex/estrous differences in an addiction-and learning implicated tissue, the hippocampus.

In WTs, we identify large divergence of gene expression between males and females only when considering single estrous stages. We further utilized gene pattern analysis to define cyclic expression patterns of the female dorsal hippocampus, and examined each pattern’s genes for enrichment in cell type markers, related transcriptomic signatures, and candidate regulators, providing an extensive resource for hypothesis generation in the study of interplay between estrous and hippocampal function. We then examined differences between *Cnih3* KOs males and KO females in specific phases of estrous and identified a profound enhancement in gene expression differences, resulting in substantially more differential genes both between the sexes.

Finally, we directly examined expression differences between WT vs KO altogether, in males, and in each estrous stage. The combined observation of broader differential expression between male and estrous stages within KOs compared to within WTs, despite the absence of substantial genotype differences, suggested that subtle changes were being induced by *Cnih3* KO to accentuate sex-differential expression. Indeed, we observed that the magnitude of sex-differential expression at diestrus or estrus was greater in the KO regardless of false discovery rate (FDR) level, supporting the hypothesis that *Cnih3* buffers against excess gene-regulatory responses to cycling sex hormones.

## Methods

### Animals, estrus staging and dorsal hippocampal dissections

Procedures were approved by the Institutional Animal Care and Use Committee at [Author University redacted until double blind review is complete]. Adult (age range 12 - 24 weeks) WT and *Cnih3* KO littermates on a C57/BL6j background were used. Genotyping was performed according to methods used by the group publishing the mouse line (Frye et al., 2021). Mice were kept in climate-controlled facilities with a 12-hour light/dark cycle and *ad libitum* access to food and water. The estrous cycle was monitored by vaginal lavage of sterile saline for at least two consecutive days prior to tissue harvesting. Samples were allowed to dry on glass slides, rinsed with water, and stained with giemsa stain (Ricca Chemical) to improve contrast and differentiate between cell types. Vaginal cell cytological analysis was used to identify estrous cycle stage by three independent observers. Estrus (E, Est) was characterized by the presence of cornified epithelial cells, metestrus (M, Met) by a mix of leukocytes and both nucleated and cornified epithelial cells, diestrus (D, Di) by leukocytes, and proestrus (P, Pro) by nucleated epithelial cells. Males and females at each stage of estrous (n= 4-7 per group) were decapitated and dorsal hippocampi were rapidly dissected over ice, tissue was snap frozen on dry ice and stored at −80 °C until use.

### Tissue processing and RNA purification

Hippocampi were lysed in 500 μL of buffer (50 mM Tris, pH 7.4, 100 mM NaCl, 1% NP-40, supplemented with RNAsin and protease inhibitors) on ice, centrifuged at 2,000 × g for 15 minutes at 4 □, and 133 μL of the supernatant was taken forward for RNA purification. This was mixed with 67 μL of Promega’s simplyRNA Tissue Kit Homogenization buffer with 1-thioglycerol (20 μL per mL), then 200 μL of Promega’s lysis, and extracted using a Maxwell RSC 48 robot (Promega) following the manufacturer’s instructions for the above Kit.

### RNAseq library preparation and sequencing

Total RNA integrity was determined using Agilent Bioanalyzer or 4200 Tapestation. Library preparation was performed with 10 ng of total RNA with a Bioanalyzer RIN score greater than 8.0. Double stranded complementary DNA (dscDNA) was prepared using the SMARTer Ultra Low RNA kit for Illumina Sequencing (Takara-Clontech) per manufacturer’s protocol. cDNA was fragmented using a Covaris E220 sonicator using peak incident power 18, duty factor 20%, cycles per burst 50 for 120 seconds. cDNA was blunt ended, had an A base added to the 3′ ends, and then had Illumina sequencing adapters ligated to the ends. Ligated fragments were then amplified for 12-15 cycles using primers incorporating unique dual index tags. Fragments were sequenced on an Illumina NovaSeq-6000 using paired end reads extending 150 bases.

### Data Analysis

RNA-seq reads were aligned to the Ensembl release 101 primary assembly (GRCm38.101) with STAR version 2.7.9a. Gene counts were derived from the number of uniquely aligned unambiguous reads by Subread:featureCount version 2.0.3. All gene counts were then imported into the R/Bioconductor package EdgeR and TMM normalization size factors were calculated to adjust for samples for differences in library size. Ribosomal genes and genes not expressed in at least ten samples greater than one count-per-million were excluded from further analysis. The TMM size factors and the matrix of counts were then imported into the R/Bioconductor package Limma. Weighted likelihoods based on the observed mean-variance relationship of every gene and sample were then calculated for all samples and the count matrix was transformed to moderated log2 counts-per-million with Limma’s voomWithQualityWeights. Differential expression analysis was then performed to analyze for differences between pairs of conditions; results were filtered to genes with Benjamini-Hochberg FDR adjusted p-values less than or equal to 0.05 except where noted. All differential expression analyses used a single input dataset and model covering all WT and *Cnih3* KO samples: *expression* ~ 0 + *group* (where group = genotype and estrous stage or male). Estrous-stage agnostic and sex-agnositc (global genotype effect) contrasts therefore collapsed contrast coefficients each stage together; for example, *WT female vs WT male DE* = *(0.25*metestrus + 0.25*diestrus …)* - *male*.

For gene pattern analysis, the degPatterns function of R package DEGreports package was used. The moderated log2 counts per million (CPM) expression values for the union of genes differentially expressed with a differential expression nominal p-value or FDR below a given threshold–noted throughout the text where these analyses were performed–across all samples from the conditions in the set of relevant pairwise comparisons was input. For example, WT estrous gene expression patterns used the union of genes DE at an FDR < 0.01 between any two stages (six total comparisons comprising four estrous stages); therefore, the log2CPM for each of the filter-passing genes from all WT female samples was inputted for clustering based on pseudotemporal expression pattern. degPatterns parameters utilized were *minc*=5 (minimum number of genes fitting a pattern to report the pattern is 5) and *reduce*=TRUE (do not leave outlier genes in the final clusters).

For ontology, cell type, and regulatory enrichments, the Enrichr web tool (Chen et al., 2013) was used, entering the list of genes derived from a differential expression analysis directly (at FDR<0.05) or those DEGs constituting a cluster per the analyses above. Result tables were downloaded and collated for databases showing putatively brain-relevant phenomenon with a reported q-value < 0.05 for the top enrichment, and at least one putatively brain-relevant, q-value significant enrichment being driven by at least 5 of the input genes. Result tables were then collated across databases in R (see *Code Availability*).

## Results

**Figure 1.**
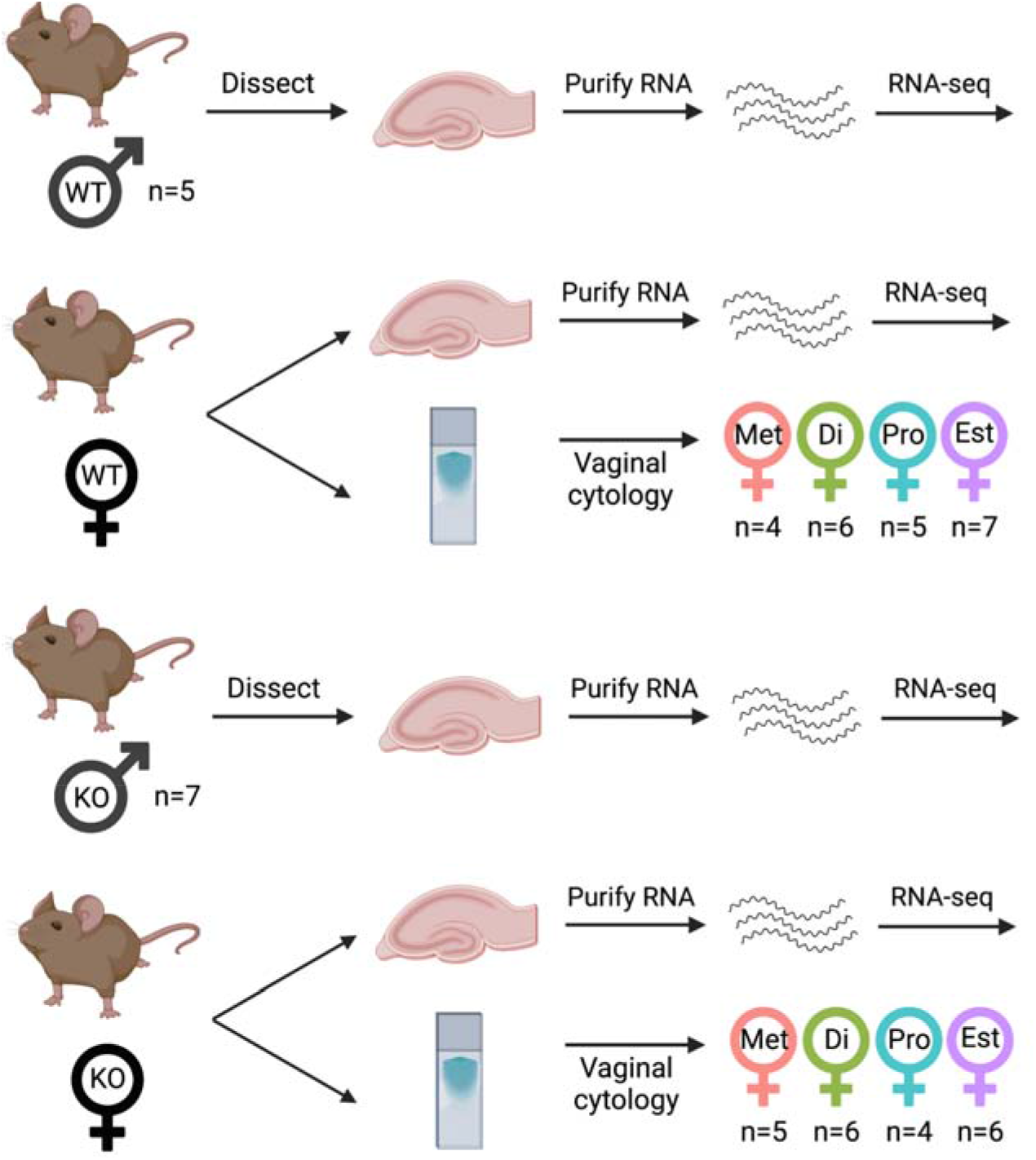
Experimental workflow. Dorsal hippocampi from male and female adult C57/B6 wild-type (WT) and *Cnih3* knockout (KO) mice were collected for RNA-sequencing (one hippocampus = one sample). Vaginal cytology was performed at time of tissue collection. Estrous stage was independently determined from cytology by 2-3 scorers. Final *n* for each genotype and estrous stage group are indicated in the figure. Met = metestrus; Di = diestrus; Pro = proestrus; est = estrus.

### Wild-type hippocampal gene expression: sex and estrous differences

#### Gene expression between wild-type male and female bulk hippocampus does not substantially differ

We first tested the 16,168 genes in our wild-type (WT) hippocampal RNA-seq data for female gene expression differences from male, examining both estrous-naïve and stage-specific differences in gene expression (**Figure 2, Supplementary Table 2a**). At the level of sex alone, we only detected 6 differentially expressed genes (DEGs) (FDR < 0.05), all from the sex chromosomes (**Fig 2A**). Examining nominally significant DEGs with a log fold-change (FC) exceeding 1.5 revealed 4 additional genes, including *Depp1* (female-upregulated, logFC 2.3, p<4×10^−3^) and *Avpr1b* (female-upregulated, logFC 2.13, p<2×10^−3^). In all, the “net” female hippocampal transcriptome did not diverge appreciably from male.

**Figure 2.**
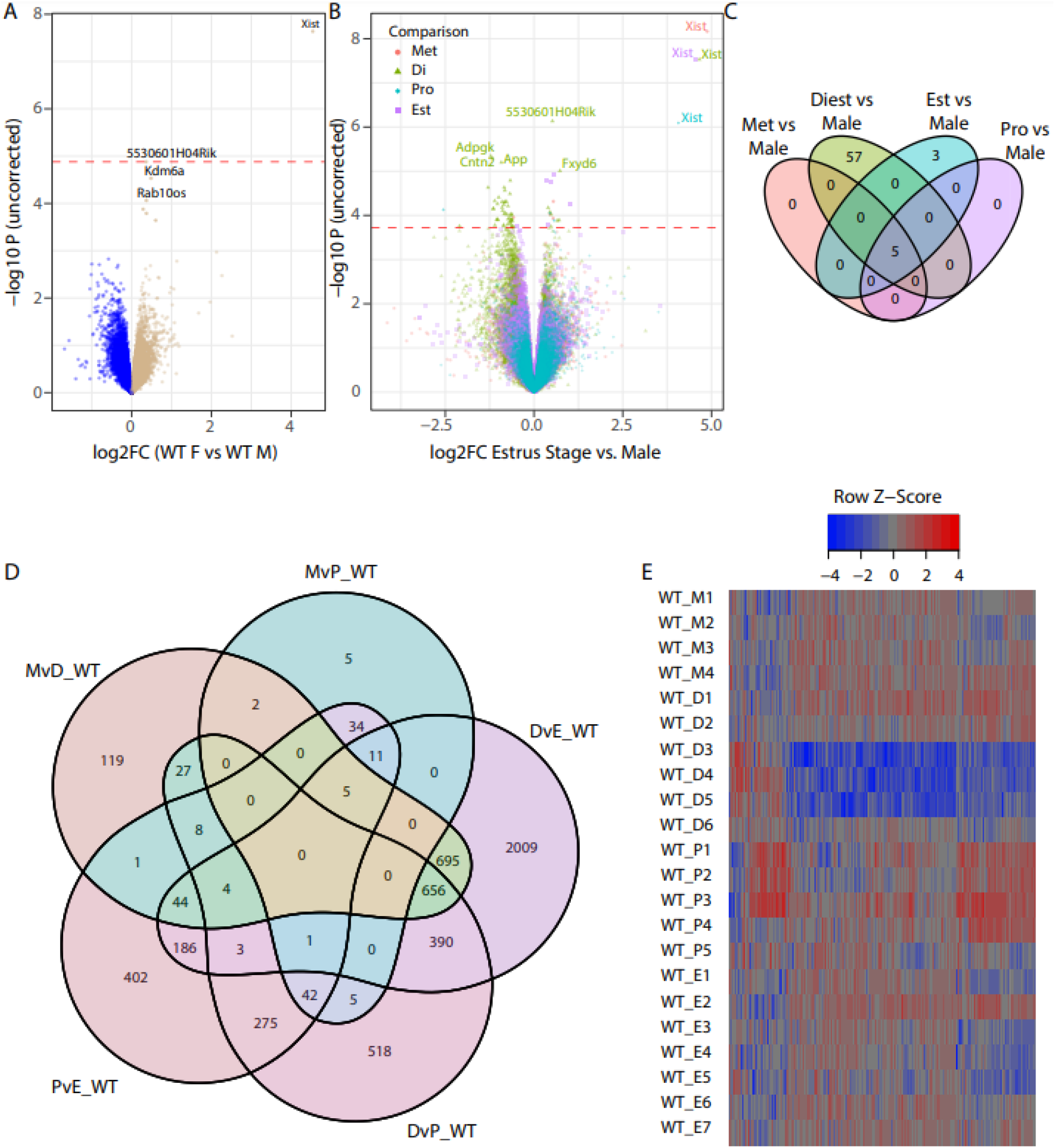
Wild-type dorsal hippocampal transcriptome varies across estrous stages. **A)** Volcano plot illustrating the effective absence of sex-differential genes when comparing wild-type males to females combined across all stages of the estrous cycle. Female enriched genes are positive. **B)** A joint volcano of differential expression analysis results for each estrous stage compared to males. The largest magnitude differences in gene expression are between diestrus and males. *Xist* is shown on the plot to illustrate the comparative magnitude of sex-specific gene expression from the sex chromosomes, Y-chromosome genes are excluded for scale. **C)** Venn diagram illustrating the number of significant DEGs between males and females for each estrous stage. The majority of differential expression occurs in diestrus. **D)** A Venn diagram of differentially expressed genes across pairwise comparisons of estrous stages, illustrating the dependence of the female hippocampal transcriptome on estrous stage. Once again, diestrus is the most distinctive of the four stages. **E)** Heat map of samplewise expression of the same genes as in panel D. Hippocampal gene expression changes across estrous stages are individually subtle but extensive in terms of the number of genes involved.

As prior experiments with *Cnih3* have illustrated, male-female differences can be influenced by the stage of the estrous cycle (Frye et al., 2021). Therefore, we performed comparisons between WT male and WT females for each estrous stage separately, identifying 65 unique genes as significant across the four comparisons (**Fig 2B-C, Supplementary Table 2b-2e**). For metestrus (Met) and proestrus (Pro), and most surprisingly, estrus (Est), we only identified sex chromosomal genes as DEGs (FDR <0.05) when compared to male. In contrast, we identified 62 significant DEGs in diestrus (Di) compared to male, 57 of which were unique to this stage (8 female-upregulated, 49 male-upregulated; these did not include *Cnih3*, consistent with qPCR reports (Frye et al., 2021)). Altogether, these findings suggest that the female dorsal hippocampus only diverges to any appreciable extent from males during diestrus.

#### Gene expression changes substantially across estrous stages within WT females

We next examined the female samples alone, comparing each estrous stage to all others to identify genes significantly fluctuating over the course of the estrous cycle. Strikingly, we identified over 5,000 unique DEGs between at least one pair of stages of the 6 combinations possible (**Fig2D-E, Supplementary Table 2f-2k**). There were no significant DEGs between Met and Est, while the other 5 pairwise comparisons yielded 105-4004 significant DEGs each.

Altogether, these findings suggest that male dorsal hippocampal gene expression diverges neither from females overall, nor from individual estrous stages. Instead, most variability in expression is constrained to females in an estrous stage-dependent manner. Diestrus appears to correspond to the most distinctive transcriptomic state of the dorsal hippocampus, in that it is the most distinct from both males and other estrous stages (in terms of DEG) at respectively modest and sprawling scales.

We confirmed our findings of estrous DEGs by comparing our directions of effect with those of significant DEGs with a high logFC from a prior study of mouse hippocampus (DiCarlo et al., 2017). For the two largest gene sets from their data also represented in our data (proestrus > estrus; diestrus < proestrus), 100% of our effect directions were in agreement (**Supplementary Figure 1**).

#### Gene expression patterns in WT hippocampus across the estrous cycle

Given the large number of pairwise DEGs identified between different stages of the estrous cycle, we were well-powered to cluster these genes by their rise and fall across the cycle to predict putative biological functions subject to estrous influence in the hippocampus. Characterization of these patterns resulted in a resource for hypothesis generation concerning sex differences in the hippocampus, including as a comparator in the study of estrous stage-specific phenotypes like those seen in *Cnih3* KO. In addition to the provided results from stage-stage comparisons (**Supplementary Table 2f-2k**), we present a series of gestalt analyses to understand these differences below.

To clump genes by their cyclic pattern of expression, we used the *DEGReport* package’s *degPatterns* function in R 4.1.2, and clustered the union of genes significant at an FDR < 0.01 (1.7k genes total) from any estrous comparison above. We identified 5 gene expression patterns total in WT (**Figure 3A; Supplementary Table S3a**), the majority of which fell into clusters corresponding to peak expression in diestrus (cluster 4) or trough expression at diestrus (clusters 1, 2), or peak expression at proestrus (clusters 1 and 3). We next analyzed each cluster’s genes in the *Enrichr* tool (Chen et al., 2013). *Enrichr* annotations of note for four of these clusters (1-4) highlighted several pertinent aspects of brain and hormonal biology (**Supplementary Table S3b**). WT estrous cluster 1, characterized by peak expression in proestrus and trough expression in diestrus (**Fig. 3A**), was enriched for oligodendrocyte marker genes (**Figure 3B**). (Indeed, 4/6 of the stagewise comparisons above showed differential expression of *Gal3st1*, whose protein product sulfonates carbohydrates in sphingolipids to produce sulfatide, a major component of myelin). Cluster 2, also characterized by trough expression in diestrus but with estrus-metestrus peak expression, showed enrichment for genes in “early estrogen response”, calcium signaling, glutamate receptor signaling, and axon guidance, as well as strong enrichment for inhibitory interneuron subtypes, glycinergic neurons, and all classes of glia, and finally regional enrichment for the molecular layer of the dentate gyrus (**Figure 3A,C**). By contrast, WT estrous cluster 3, with peak expression across diestrus and proestrus, was enriched for protein interactors of estrogen receptor *Esr1* with only weak enrichment for dopaminergic, glycinergic, and subtype-nonspecific neuron markers (**Supplementary Table S3b)**. Finally, the genes sharply peaking in diestrus of cluster 4 were strongly enriched for *Sncg*+ neurons and hippocampal CA3 neurons from Allen Brain Atlas single-cell RNA-seq, and were enriched for genes upregulated by knockdown of *RELA, Neurod1*, or *Mecp2*, or by overexpression of *Neurog3* (**Supplementary Table S3b)**. These findings highlight higher-order biological changes in the dorsal hippocampus occurring at different stages in the estrous cycle, with especially strong cell type and regional signatures for genes following the pattern of cluster 2.

**Figure 3.**
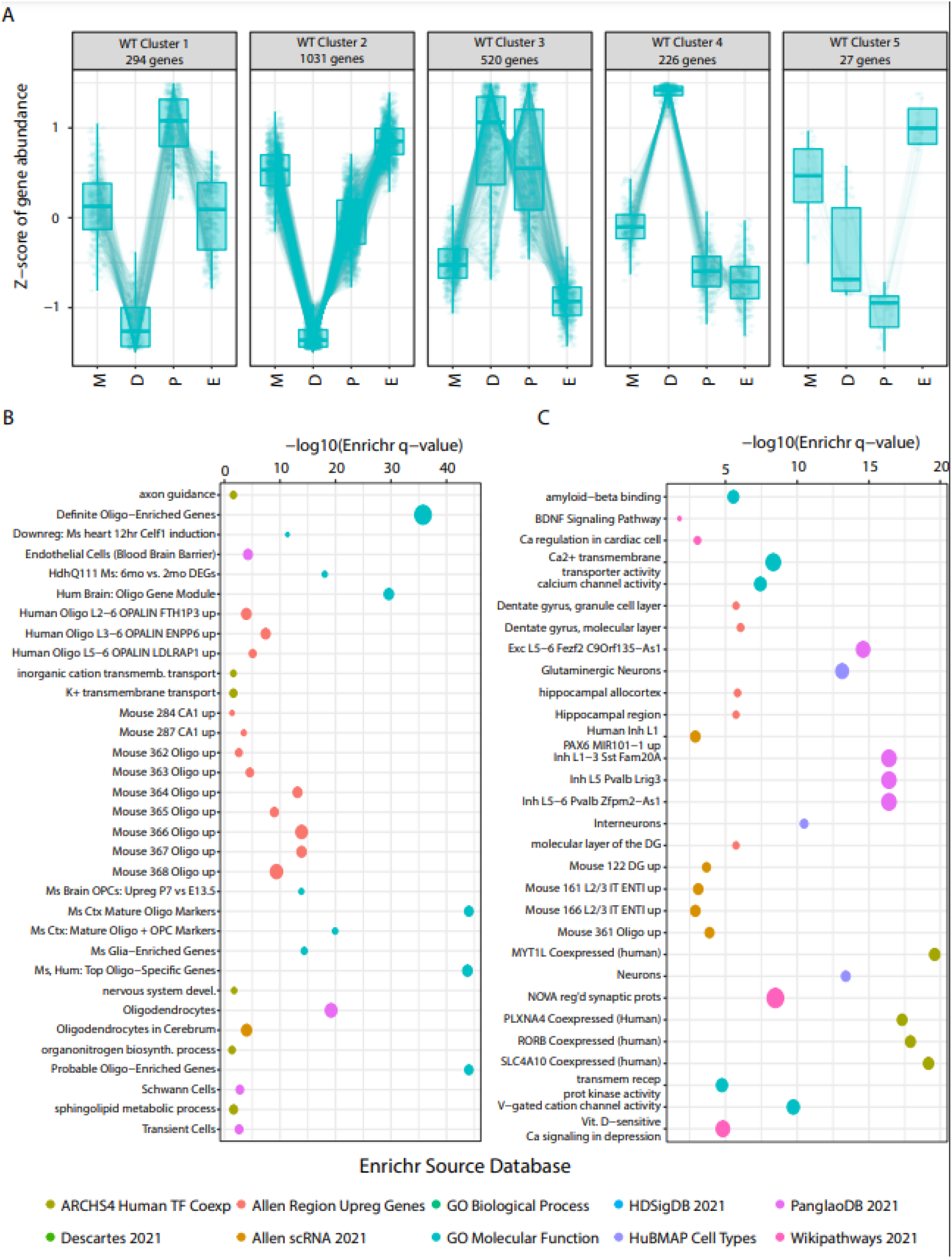
Wild-type female cyclic hippocampal gene expression patterns across estrous. **A)** Patterning analysis of gene expression for DEGs between any pair of stages at FDR <0.01 identifies five clusters of expression fluctuation across the estrous cycle. **B)** Selected Enrichr analysis results for cluster 1 from panel A. Specific terms are highlighted in the plot, all with log odds ratio (OR) of the cluster gene set > 2.5, all enrichment (uncorrected) p-values < 0.05, with at least 8 genes from the cluster included in the enriched annotation term. Point colors indicate which Enrichr dataset each term comes from, while point sizes are scaled to the log OR. **C)** Selected Enrichr analysis results for cluster 2 from panel A. All terms meet the same filtering criteria as in Panel B.

### *Cnih3* knockout effects on dorsal hippocampus

We first examined KO and WT *Cnih3* sequencing alignments to confirm that *Cnih3* exon 4 was indeed absent, as expected for this mouse line (**Supplementary Figures 2 and 3**). This mutation induces a frameshift and thus predicted loss of function in *Cnih3*. Subsequently, we examined expression of all genes for the KO mice in the same series of approaches as for WT above, which are on the whole presented in a similar structure to those above for sake of comparability. Finally, we perform KO vs WT comparisons for each sex/estrous stage, and finally, we describe the overall key patterns of transcriptomic alterations identified in the *Cnih3* KO hippocampus.

#### Gene expression differences between *Cnih3* KO males and females are subtle but far outnumber WT sex differences

*Cnih3* KO males (n=7) and females (all estrous stages considered jointly, n=21) showed much starker differential expression in the dorsal hippocampus, with 849 genes detected as differentially expressed at FDR < 0.05 (**Figure 4A, Supplementary Table 4a**). Notably, only 18.6% of these genes had an absolute logFC exceeding 0.5, suggesting the vast majority of estrous-nonspecific, sex-differential expression in the *Cnih3* KO hippocampus is subtle in nature. The two autosomal genes from this comparison with a logFC of > 1.5 were perilipin 4 (*Plin4*), and as observed at nominal significance in the WT sex comparison above, prolactin (*Prl*).

**Figure 4.**
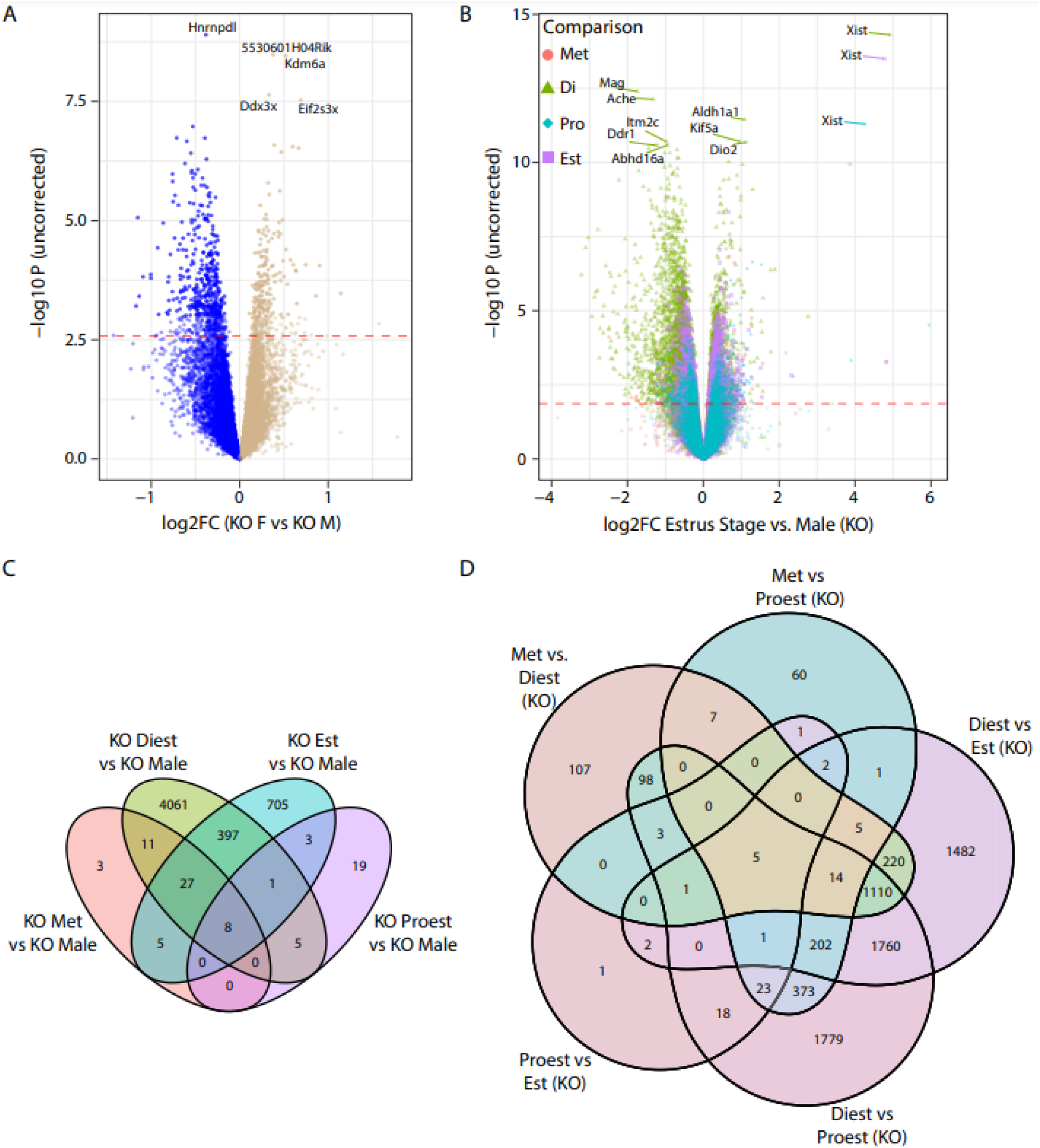
*Cnih3* knockout hippocampal transcriptome shows greater differences between sexes and estrous stages. **A)** At the level of males compared to all females, the *Cnih3* knockout line shows a substantial increase in the number of significant DEGs compared to WT (Figure 2A). **B)** A joint volcano of differential expression analysis results for each KO estrous stage compared to KO males. The largest magnitude differences in gene expression are between diestrus and males, consistent with the WT pattern of diestrus constituting the most distinct transcriptional state. **C)** Likewise, the greatest number of DEGs between males and an estrous stage are found at diestrus, as shown in the Venn diagram of significant genes from the comparisons in panel B. **D)** Diestrus is likewise the most distinctive transcriptional state within *Cnih3* KO females, illustrated in a Venn of genes significantly DE between any two stages, where the preponderance of stage-pair-specific DEGs correspond to a diestrus comparison.

Considering the estrous-stage specific sex differences in behavior and dorsal hippocampal architecture we previously observed in *Cnih3* KO mice, we also examined males compared to each stage of the estrous cycle independently (**Figure 4B, Supplementary Table 4b-4e**). Here, also, we detected a far greater number of DEGs compared to the same approach in WT: an 80-fold greater number of unique DEGs across the four KO comparisons, totaling 5245, compared to the 65 from the four contrasts in WT. The overall distribution of DE events was similar to WT, albeit on a much larger scale, with 54, 4510, 36, and 1146 DEGs for Met, Di, Pro, and Est, respectively, compared to males (**Figure 4C-D**). Likewise, the number of high-magnitude (log FC > 1.5) DE events for Di vs. males totaled 145 in KO, vs. 6 in WT. Altogether, these findings suggest that *Cnih3* knockout accentuates sex differential gene expression in both a global and estrous stage-specific manner.

#### *Cnih3* KO females essentially retain most WT estrous expression patterns outside of small sets of proestrus-stimulated genes

As in our WT data, we then performed pairwise comparisons of estrous stages in the *Cnih3* KO samples (**Supplementary Table 4f-4k**). On the whole, these differential expression sets were comparable in size, with the exception of substantial increases in the number of FDR significant DEGs between Di-Pro and Di-Est for KO relative to WT (increases of ~3400 and ~800 genes, respectively; **Figure 4E**). Despite the similar gene set *sizes*, the DEGs between the WT and KO comparisons were only about 40-50% shared (**Supplementary Table 4l**), additionally suggesting perturbations to ordinary estrous cycle gene expression.

To clarify whether the broad-scale cyclic patterns of gene expression across the estrous cycle we identified in WT above were intact in *Cnih3* KO mice, we Z-scored the KO expression levels of the same genes and overlaid them into the WT clustering space to compare their cyclic expression to WT (**Figure 5A**). While many clustering patterns seemed to be generally retained–if sometimes with larger gene-set level variance in KO–there were notable discrepancies between WT and KO cycling patterns for clusters 1 and 3, two clusters defined by genes with peak expression in proestrus.

**Figure 5.**
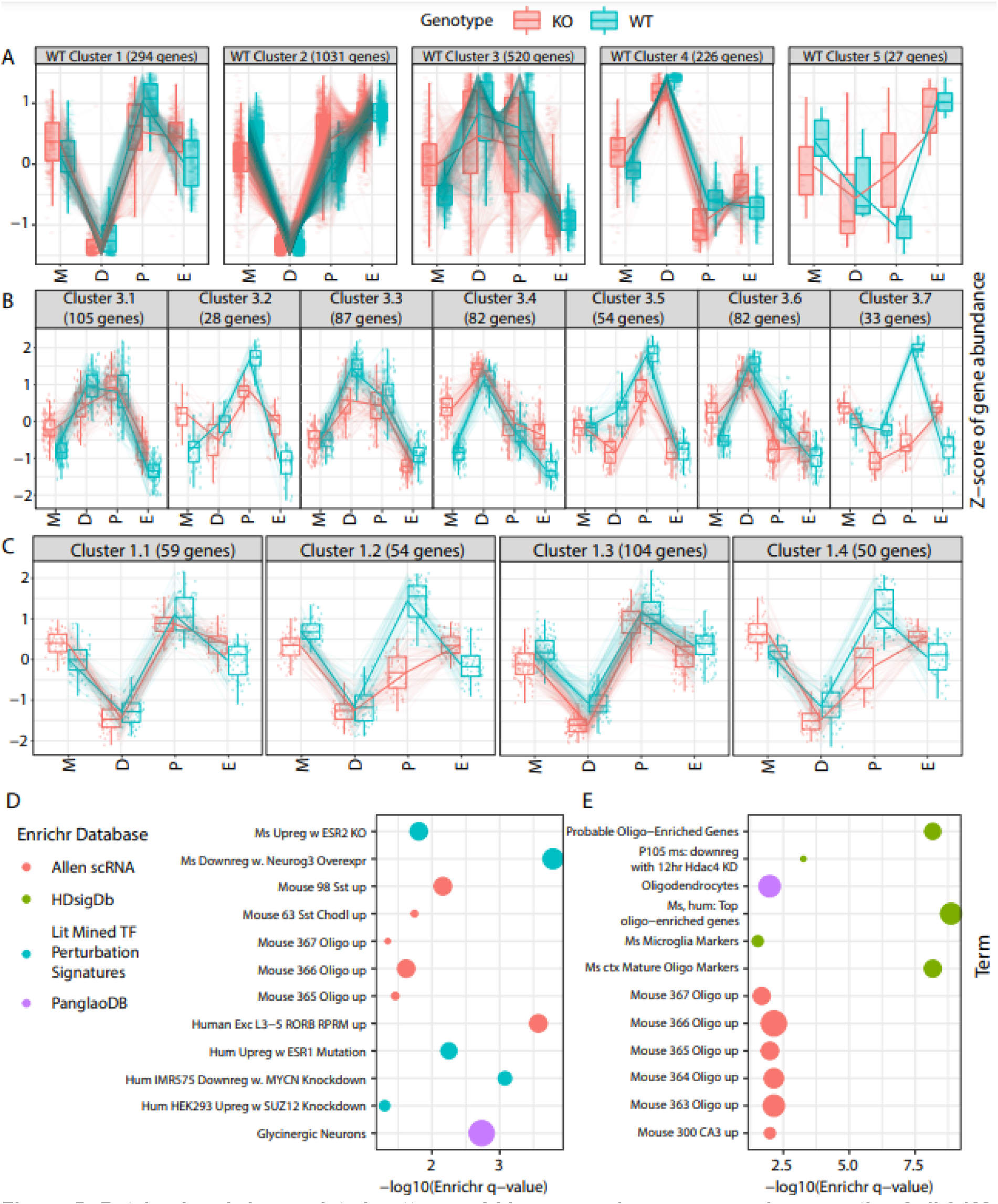
Retained and dysregulated patterns of hippocampal gene expression over the *Cnih3* KO estrous cycle. **A)** The same genes from the same five wild-type clusters of cyclic expression shown in Figure 3A are again shown, but now with the KO expression levels of those genes additionally plotted. **B)** Re-clustering of WT cluster 3 using both WT and KO data identified 7 sub-clusters (3.1, 3.2…), corresponding to a specific pattern of regulation in WT and a specific pattern in KO. Three of these clusters (3.3, 3.5, 3.7) show an attenuation of gene upregulation as the cycle progress, especially in the Met -> Di and Di -> Pro transitions. **C)** Similar re-clustering of WT cluster 1 identified 4 pattern subgroups, two of which also featured attenuation of upregulation at proestrus compared to WT (1.2, 1.4). **D)** Enrichr analysis of the genes in subclusters 3.3, 3.5, and 3.7 combined highlight that these genes are perturbed by estrogen receptor perturbation experiments and enriched in oligodendrocytes and inhibitory neuron classes. **E)** Enrichr analysis of genes in subclusters 1.2 and 1.4 combined were highly enriched for oligodendrocyte genes across several annotation sets, and implicate *Hdac4* as an upstream regulator. Filters for both Enrichr plots are the same as for Figure 3, except with a minimum term-gene set overlap of 5 rather than 8 to account for the smaller gene set sizes of these subclusters.

To further dissect the alterations to clusters 1 and 3, we re-performed clustering on the genes defining the WT cluster using KO and WT data combined, resulting in 4 and 7 subclusters of expression (*i.e*., subclusters with a specific pattern in WT and a specific pattern in KO; we call these e.g. cluster 3.1, 3.2…), respectively. This co-visualization, and the fact that unitary WT clusters divide up into multiple parts when KO data is also considered, confirmed that the KO estrous cycle patterns of certain subsets of genes were highly divergent from their WT counterparts. In the case of WT cluster 3, our combined-genotypes analysis revealed three subclusters (3.3, 3.5, and 3.7) with attenuated gene upregulation at proestrus in KO mice (**Figure 5B**). To understand functional correlates of these genes, we performed Enrichr analysis of the genes from subclusters 3.3, 3.5, and 3.7 combined, revealing oligodendrocytes and, interestingly, genes found to be upregulated in estrogen receptor knockout datasets (**Figure 5C**, **Supplementary Table 4m-4n**). Likewise, we noticed an attenuation of gene upregulation in KO mice over the diestrus-estrus stages for subclusters 1.2 and 1.4 (**Figure 5D**), whose parent cluster had shown enrichment for oligodendrocyte marker genes and potassium channels. Enrichr analysis of these two combined subclusters strikingly revealed that 52 of these 104 genes were in “Mouse Cortex Mature Oligodendrocyte And Progenitor Cell Markers” as defined in (Doyle et al., 2008); the 104 gene set was also enriched in an additional oligodendrocyte marker list similarly mined from literature (Cahoy et al., 2008), and in various combinations of oligodendrocyte genes identified by single cell RNA-sequencing (**Figure 5E, Supplementary Table 4m-4n)**. These findings very strongly suggest that the *Cnih3* KO mouse has specific deficits in oligodendrocyte gene upregulation in response to proestrus.

#### WT-KO differential expression is unremarkable within sex/estrous stage, against expectations

While above, we contrasted our within-KO comparisons to analogous WT comparisons, we examined WT-KO directly to better understand effects of the knockout on hippocampal gene expression. Contrasting all KO and WT samples, regardless of sex, identified 514 significant DEGs (35 with an absolute logFC > 0.5) (**Figure 6A**), again suggesting overall subtle effects of the *Cnih3* KO on gene expression (**Supplementary Table 5a**). *Enrichr* analysis of these 514 genes revealed highly significant overlap with genes from dozens of transcription factor perturbation experiments, including *Neurod1* knockdown and human cell culture *GATA6* overexpression. This gene set was also highly enriched for protein interactors of estrogen receptor alpha (*Esr1*). *Cnih3* KO DEGs were also enriched for neuronal markers including those of *Scng*-expressing interneurons. Intriguingly, and in direct contrast to the enrichment of KO estrous cycle-dysregulated for highly-expressed genes from oligodendrocyte single-cell RNA-seq clusters, WT-KO DEGs were instead enriched for those genes significantly *depleted* from the same oligodendrocyte clusters relative to other Allen Atlas single-cell RNAseq cell types (**Figure 6B**, **Supplementary Table 5b**).

**Figure 6.**
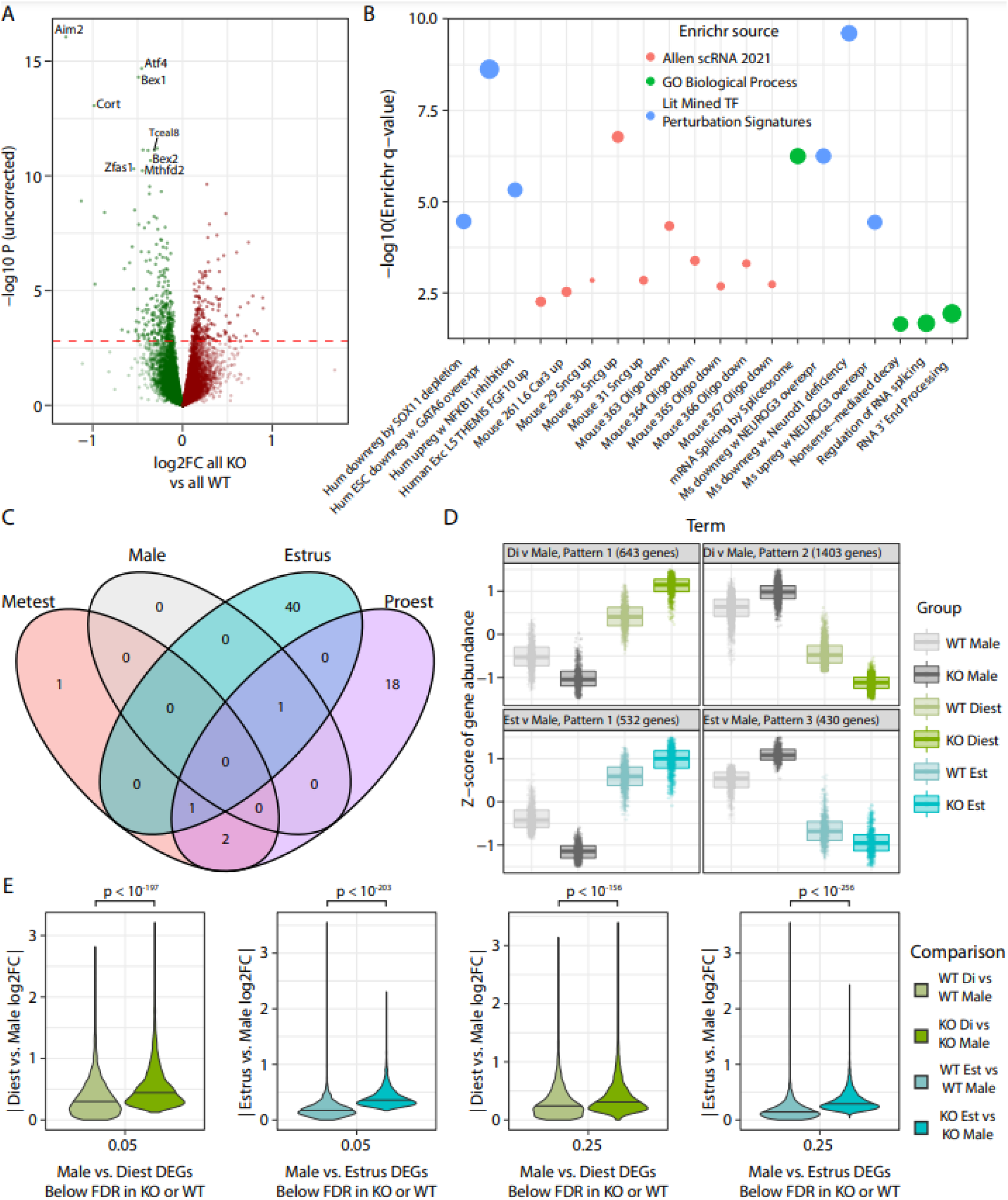
Global features of *Cnih3* KO: Splicing, synuclein, SST, subsurface neurons, and sex difference accentuations. **A)** Comparing all KOs to all WTs, regardless of sex or estrous stage, identifies hundreds of significant DEGs. **B)** Global DEGs in *Cnih3* KO are enriched for multiple neural subtypes of the mouse brain marked by synuclein gamma (*Sncg*), deep layer excitatory neurons, *Sst*-expressing interneurons, and ontology terms related to several forms of RNA processing. Notably, the same oligodendrocyte subtypes identified as *enriched* for genes dysregulated in the KO estrous cycle are *depleted* (i.e., lowly-express) in the global DEGs of the KO mouse. All terms plotted were nominally significant for enrichment and met filters for a minimum log OR of 2 and a minimum of 8 DEGs overlapping with the annotation term set. **C)** A surprisingly small number of DEGs are identified by comparing single estrous stages or males between KO and WT. (Stages not shown have 0 significant DEGs.) **D)** Analysis of genes DE between KO males and a KO estrous stage at FDR < 0.01 showed several sizable groups of genes where Cnih3 KO accentuated a normal sex difference seen in WTs. **E)** The absolute log2 fold-change for those genes DE between KO males and KO diestrus or KO males and KO estrus is shown alongside the absolute log2FCs for the respective WT comparison. At any FDR thresholds chosen, the absolute magnitude of DE is significantly (Wilcoxon test) greater in the KO comparison, indicating virtually global potentiation of sex differences in gene expression.

#### *Cnih3* KO results in accentuated sex-differential expression compared to WT, especially in diestrus

Subsequently, we performed within-sex/estrous stage genotype comparisons. Given the extremely large increase in number of sex- and male-estrous DEGs within KOs, we expected to see comparable numbers of genotype-differential genes within each group. Shockingly, however, only 1-42 FDR-significant DEGs were identified in the genotype comparisons (**Supplementary Table 5c-g**), totaling 63 unique genes (**Figure 6C**), with the most differences seen between WT and KO proestrus. Surprisingly, we observed only 1 DEG between WT and KO diestrus, despite this stage being responsible for the most DEGs between male and other estrous stages in both WT and KO mice.

Given the simultaneous finding of increased sex/estrous-differential gene expression within KOs despite little significant difference within any sex/estrous group across genotypes, we reasoned that there must be relatively subtle fluctuations occurring between WT and KO within each group such that a) only small changes occur within multiple subgroups of KO relative to their sex/estrous-matched WT counterparts (yielding few significant DEGs in these comparisons), while simultaneously, b) the magnitude of these changes is in opposing directions between males and one or more estrous stages, resulting in a greater number of DEGs identified in KO male-stage comparisons. To examine whether this hypothesis was valid at the level of KO male-vs-stage DEG sets, we again utilized the *degPattern* algorithm, now with the intention of visualizing changes from WT in expression patterns of genes identified as DE in KO male-stage comparisons. Indeed, for the two KO estrous stages with over 1k significant DEGs compared to KO male, we see that most DEGs are altered in opposite directions in male and female KOs, resulting in accentuated sex differences across large portions of the hippocampal transcriptome (the two patterns from each comparison showing this effect are in **Figure 6D**; all patterns are in **Supplemental Tables 6A-6B**). Enrichr analyses of genes following each pattern for these two male-stage comparisons (such as M>diestrus in WT with male upregulation and female downregulation in KO) are provided in **Supplemental Table 6C**, and recurring terms across genes with different patterns are tabulated in **Supplemental Table 6D**. To examine this phenomenon at a global scale, we also took all genes differentially expressed between KO males and KO diestrus or KO estrus at varying levels of FDR stringency, and plotted their absolute log2 fold change values in that comparison for WT or for KO mice (**Figure 6E**). At all FDR thresholds tested, a non-parametric Wilcoxon test identified extremely significant positive deviation of the KO log2 fold changes from those seen in WT, confirming a general net increase in sex-differential expression between Cnih3 KO males and KO females in diestrus or estrus.

## Discussion

Overall, driven by the following considerations A) sex differences in psychiatric disorders, including addiction; B) the role of sex hormones in regulating the HPA axis; C) the role of stress in drug reinstatement (Sherry A. McKee, 2015); D) the molecular and behavioral integration of both sex and stress hormone signals in the hippocampus (Bao et al., 2006; Frick et al., 2015b; Joshua A Gordon, 2016); E) the prior work showing the influence of estrous stage on hippocampal physiology and spatial learning (Frye, 1995; Warren and Juraska, 1997); and F) the sex and estrous specific differences in hippocampal phenotypes in *Cnih3* KO females, we conducted a well powered study to understand the transcriptional effects of estrous on dorsal hippocampal gene expression. We found that on average, the brain is well buffered against sex differences in expression, with the average gene expression showing few differences between males and females. However, at specific stages of the estrous cycle, females differed more substantially from males. Likewise, though less extensive, estrous stages also showed differential expression between one another; in comparing single stages against one another or to males, diestrus was consistently the most transcriptionally distinct state from males and from across the estrous cycle.

We examined data from wildtype females and identified various changes across estrous cycle phases that may have interesting biological relevance to estrous cycle specific changes in hippocampal physiology and behavior. For genes in Cluster 1, with trough expression in diestrus and peak expression in proestrus - in which estradiol goes from lowest to highest, and progesterone starts high and begins to decrease - we observed enrichment in oligodendrocyte markers. Myelin, which is made by oligodendrocytes and increases efficiency of synaptic transmission, has been shown to increase after increased neuronal activity in the motor cortex (Gibson et al., 2014), and this was necessary for the motor function enhancement seen in their system. It is interesting to note that LTP, which is characterized by increased neuronal transmission, is enhanced during proestrus (Warren et al., 1995) with some reporting improved object-based spatial learning during this phase (Frye, 1995). Thus, it is possible that changes in myelination might help support this increased LTP and learning. For genes in Cluster 2, with peak expression in estrus/metestrus and trough expression in diestrus - in which estradiol is high in estrus then fluctuates and progesterone starts low and begins to increase - we observed enrichment in genes involved in calcium and glutamate receptor signaling. Previous research has shown that estradiol improves recognition and spatial learning, and increases hippocampal spine density in CA1 (Woolley and McEwen, 1993) which requires calcium and glutamate receptor signaling. Thus, it will be interesting to investigate whether some of the receptors upregulated here mediate the improved spatial learning seen at this phase.

We then examined data from *Cnih3* knockouts in the same manner, identifying a much more marked extent of sex-differential expression when considering clumping all estrous stages together, and likewise between males and single stages. Using our WT estrous cycle gene expression patterns-and given the estrous-stage specific behavioral changes previously observed in *Cnih3* KO mice (Frye et al., 2021)–we examined whether KO expression patterns deviated from wild-type expression over the estrous cycle. We indeed identified specific subsets of genes with blunted upregulation in KO over estrogenic stages of the cycle, especially proestrus. These dysregulated gene subsets implicated oligodendrocytes, glycinergic neurons, and somatostatin (SST) interneurons potentially involved cell types, downstream of myriad candidate regulators including *Hdac4*, *Klf4*, *Neurog3*, and the estrogen receptor *Esr1*.

*Cnih3* was of interest because human genetic association studies suggested polymorphism in this region could be protective against opioid dependence (Nelson et al., 2016), a disease that involves hijacking of normal reward mechanisms including learning and memory processes. Cnih3 is annotated as an AMPA receptor trafficking protein, initially based on sequence homology. It was later shown to bind AMPA receptors and alter gating properties in culture systems (Coombs et al., 2012; Shanks et al., 2014). A subsequent focusing on phenotypes accessible to hippocampal slice physiology had shown little function of *Cnih3*, except in the context of co-deletion of *Cnih2*. When both proteins were deleted, physiological studies in the acute slices revealed a phenocopy of several aspects of GluA1 KO (a subunit of AMPA channels), including altered mEPSC amplitude and kinetics, and deficits in long term potentiation (Herring et al., 2013). However, behavioral effects were not assessed, and notably these slices were all generated from pre-pubescent animals as is standard in the field. Finally, a recent investigation of hippocampal learning and memory function via Barnes Maze in *Cnih3* KO mice revealed no main effects of genotype initially, but a surprising amount of variance in females. Subsequent exploration of this led to the discovery of estrous-stage specific effects of *Cnih3* levels both in KOs, as well as in hippocampal specific *Cnih3* overexpression (Frye et al., 2021). Furthermore, synaptic physiology (in adult slices) as well as biochemical and immunofluorescent analyses of synapses revealed against stage specific alterations in hippocampal properties across multiple levels. All of these results led to the hypothesis that Cnih3 in some way buffers against hormone dependent sex differences, with the loss of the protein unmasking deficits in KO females.

We therefore directly examined expression differences in WT vs KO altogether and between males or estrous stages. The 514 genes we identified as differentially expressed between genotypes highlighted some shared enrichments with estrous-(dys)regulated gene sets at the level of cell types (SST interneurons, *Scng*-expressing neurons) and transcriptional regulators (*Neurog3*); however, most functional enrichments were distinct, spanning several forms of RNA processing and transport, as was the case for candidate upstream regulators, which included *Neurod1, Hsp90*, and, interestingly, X-binding protein 1 (*Xbp1*). When we compared single estrous stages or males across genotypes, however, we identified very little differential expression. The combined observation of broader differential expression between male and estrous stages within KOs compared to within WTs, despite the absence of genotype differences, suggested to us that subtle changes were being induced by *Cnih3* KO to accentuate sex-differential expression. Indeed, we observed that the magnitude of sex-differential expression at diestrus or estrus was greater in the KO regardless of FDR level, confirming our hypothesis that *Cnih3* buffers against excess gene-regulatory responses to cycling sex hormones.

Altogether, we deeply characterize hippocampal gene expression patterns over estrous in WT mice, characterize the *Cnih3* KO hippocampal transcriptome, and identify a surprising potentiation of sex differential gene expression in this knockout line. The data and supplements from these analyses provide extensive gene annotations for WT regulatory patterns, their dysregulation in *Cnih3* KO, and a well-powered dataset illustrating the role of estrous stage in defining sex-differential gene expression. These analyses and data provide an extensive resource for the study of sex- and estrous-differential gene expression in the mouse hippocampus.

## Data Availability

Data will be made available on GEO upon publication.

## Code Availability

Code for read QC, alignment, and generation, and filtering of count data is available from the authors by request. The raw unfiltered counts, a filtered count matrix as used for analysis, code to analyze the filtered counts in limma, and scripts for all analyses/plotting thereafter (with the exception of Enrichr analyses themselves, which were executed and collected through the Enrichr web tool) will be made available on Bitbucket at publication, or by reasonable request in personal correspondence to BM.

## Acknowledgements

The authors would like to thank Lexi Harris, Kristy Begmann, and Jessica Higginbotham for their assistance. This work was supported by the NIH (R33DA041883) and the Simons Foundation (734069).

---

**Supplementary Tables**

**Supplementary Table 1: RNA-seq QC results.**

**Supplementary Table 2a-2k: Wild-type differential expression contrast results.** Sheets are named according to the contrast performed and cover all genes analyzed as described in methods.

**Supplementary Table 3a-3b: Wild-type estrous gene cycling patterns and pattern annotations. A)** Pattern assignment of genes DE between any two estrous stages in WT at an FDR < 0.01. **B)** Enrichr analysis results for each gene set defined in sheet A, with all enrichments achieving at least nominal significance. Results for each gene set only are reported for Enrichr databases with nominally significant enrichment of at least one term of potential nervous system or endocrine implications.

**Supplementary Table 4a-4n: *Cnih3* KO mouse differential expression contrast results, number of DEGs and degree of DEG overlap between KO and corresponding WT contrasts, and re-clustering assignments of estrous cycling genes from WT clusters 1 and 3. A-K)** Sheets are named according to the contrast performed and cover all genes analyzed as described in methods. **L)** Table summarizing estrous stage-stage comparisons in WT and KO by number of total DEGs in each genotype and number of shared DEGs across the two genotypes for each stagewise comparison. **M)** Gene clustering reassignment of those genes from WT clusters 1 and 3 when also including KO data as a second set of data points. *Orig.WT.Clust* indicates the parent cluster (i.e. from **Supplementary Table 3A**), while *joint.WT.KO.subcluster* indicates the subcluster that resulted (i.e. as shown in **Figure 5C-5D**). These are the same indexing values used to describe subclusters in the main text/figures (e.g., subcluster 1.3 has column values of Orig.Wt.Clust as 1, joint.WT.KO.subcluster as 3). **N)** Enrichr results for the union of genes in subclusters 1.2 and 1.4 or in the union of subclusters 3.3, 3.5, and 3.7. Results for each gene set only are reported for Enrichr databases with nominally significant enrichment of at least one term of potential nervous system or endocrine implications.

**Supplementary Table 5a-5g: Genotype differential expression contrast results. 5A,5C-5G)** Sheets are named according to the contrast performed and cover all genes analyzed as described in methods. **5B)** Nominally significant Enrichr results for genes DE between all KOs and all WTs considered together.

**Supplementary Table 6a-6d: Patterns of altered sex-differential expression between WT and *Cnih3* KOs for diestrus and estrus. A)** Cluster assignment of KO male vs diestrus DEGs for expression patterns across WT male, KO male, WT diestrus, and KO diestrus. Two such patterns are shown in **Figure 6D**. **B)** Cluster assignment of KO male vs estrus DEGs for expression patterns across WT male, KO male, WT diestrus, and KO diestrus. Two such patterns are shown in **Figure 6D**. **C)** Enrichr annotations for the diestrus-male KO DEGs with KO/WT sex DE patterns 1 or 2 as listed in sheet 6A. **D)** Enrichr annotations for the diestrus-male KO DEGs with KO/WT sex DE patterns 1 or 2 as listed in sheet 6A.

**Supplementary Table 7a-7b: Samplewise expression data for filter-passing genes and metadata for analysis. A)** Moderated log2 counts per million (CPM) values for all samples analyzed in the study. These values are the values used for generation of plots using expression/Z-scored expression (e.g., expression clusters). Column names are annotated in sheet **B)** metadata (genotype, sex, estrous stage).

**Supplementary Figure 1.**
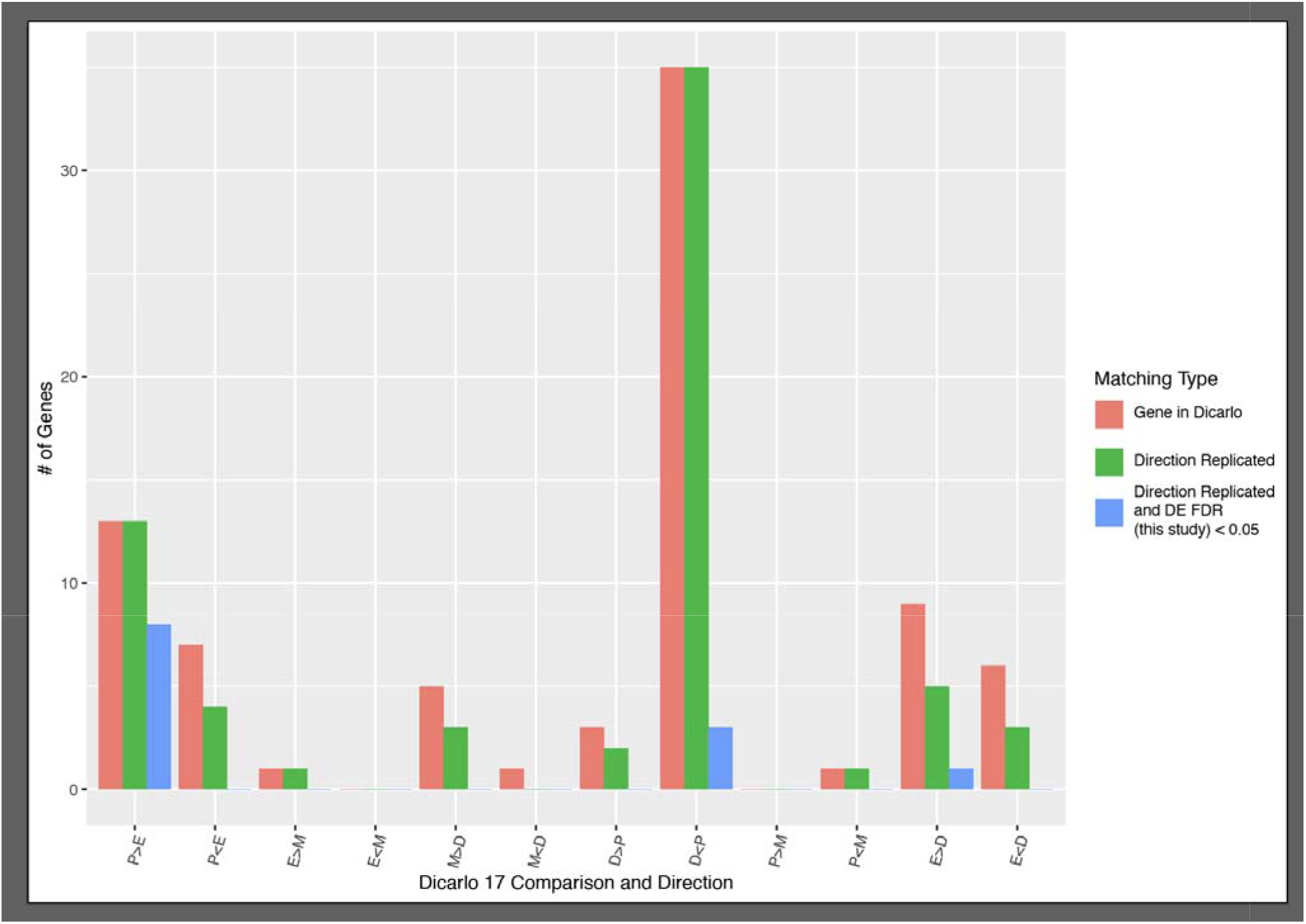
Replication of WT estrous stage pair DEGs between the present study and DiCarlo 2017. Each X-axis value represents one direction of expression change between the first and second listed stages (*e.g*., P>E signifies genes upregulated in proestrus relative to estrus; P<E signifies downregulation in proestrus relative to estrus). All 6 comparisons (12 directional changes) were examined in WT both here and in Dicarlo 2017. M; metestrus; D: diestrus; P: proestrus; E: estrus.

**Supplementary Figure 2.**
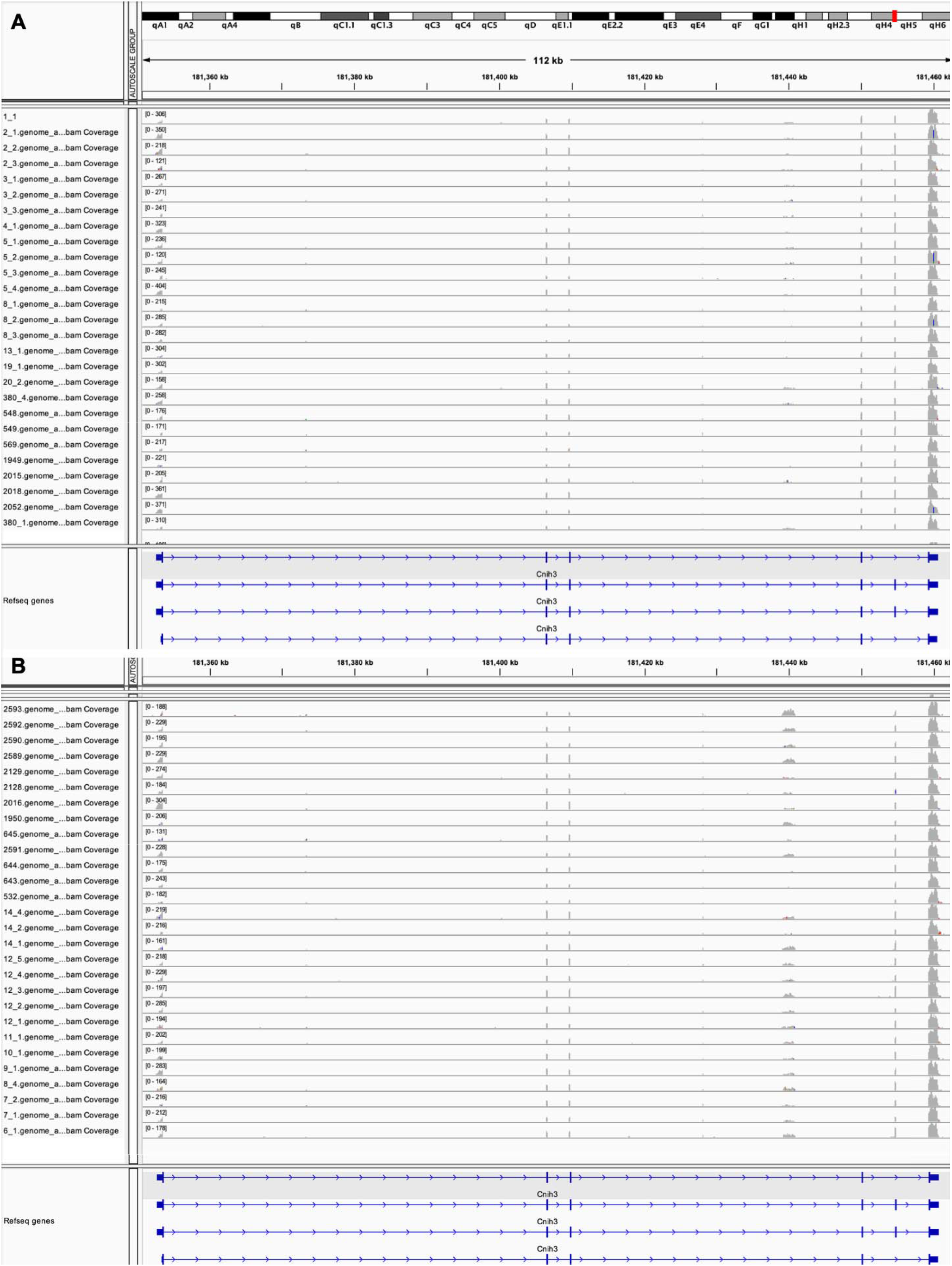
*Cnih3* RNA-seq read coverage in each analyzed sample. **A)** Coverage in WT samples. **B)** Coverage in KO samples.

**Supplementary Figure 3.**
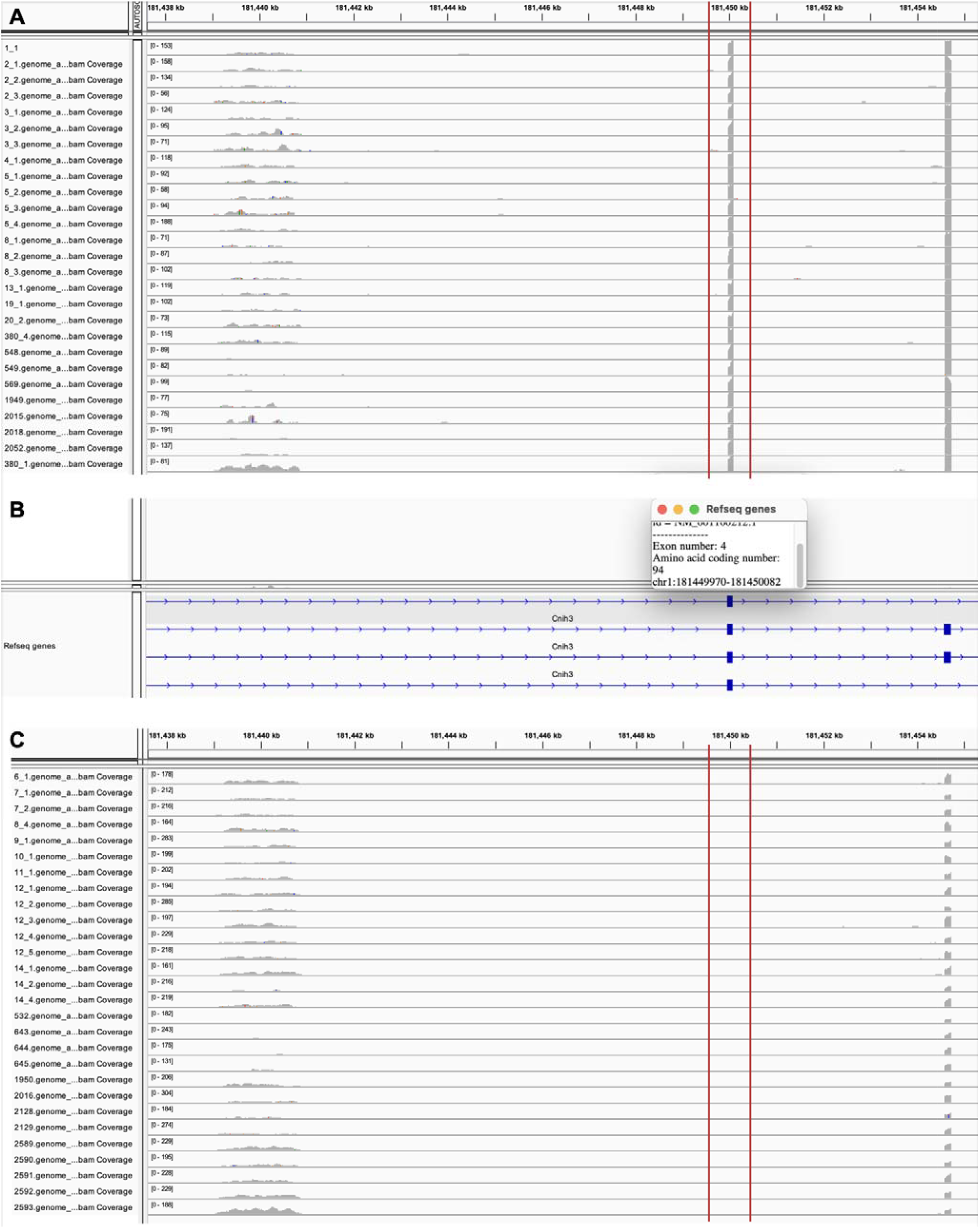
*Cnih3* read coverage recapitulates the KO strain loss of exon 4 previously reported. Higher zoom of the exon 4 region of *Cnih3* is shown for **A)** WT samples and **C)** KO samples, with the Cnih3 gene track and corresponding information on exon 4 shown between (**B**).

